# Spike-and-wave discharges of absence seizures in a sleep waves-constrained corticothalamic model

**DOI:** 10.1101/2022.10.31.514510

**Authors:** Martynas Dervinis, Vincenzo Crunelli

**Affiliations:** Neuroscience Division, School of Bioscience, Cardiff University, Museum Avenue, Cardiff CF10 3AX, UK

**Keywords:** cortex, thala mus, T-type Ca^2+^ channels, GABA-A inhibition, GABA-B inhibition

## Abstract

**Aims:** Recurrent network activity in corticothalamic circuits generates physiological and pathological EEG waves. Many computer models have simulated spike-and-wave discharges (SWDs), the EEG hallmark of absence seizures (ASs). However, these models either provided detailed simulated activity only in a selected territory (i.e. cortical or thalamic) or did not test whether their corticothalamic networks could reproduce the physiological activities that are generated by these circuits.

**Methods:** Using a biophysical large-scale corticothalamic model that reproduces the full extent of EEG sleep waves, including sleep spindles, delta and slow (<0.1 Hz) waves, here we investigated how single abnormalities in voltage- or transmitter-gated channels in the neocortex or thalamus lead to SWDs.

**Results:** We found that a selective increase in the tonic γ-aminobutyric acid type A receptor (GABA-A) inhibition of first order thalamocortical (TC) neurons or a selective decrease in cortical phasic GABA-A inhibition are sufficient to generate ∼4 Hz SWDs (as in humans) that invariably start in neocortical territories. Decreasing the leak conductance of higher order TC neurons leads to ∼7 Hz SWDs (as in rodent models) while maintaining sleep spindles at 7-14 Hz.

**Conclusion:** By challenging key features of current mechanistic views, this simulated ictal corticothalamic activity provides novel understanding of ASs and makes key testable predictions.

## INTRODUCTION

Absence seizures (ASs) are genetic, generalized, non-convulsive seizures characterized by sudden, relatively brief lapses of consciousness that are invariably accompanied by spike-and-waves-discharges (SWDs) in the EEG.^1-5^ It is well-established that both the clinical and the electrographic symptoms of ASs originate from aberrant activity of corticothalamic networks^5^ and a number of genetic abnormalities have been identified in humans with ASs.^6^ However, our current understanding of how these genetic deficits lead to the ictal EEG activity observed during ASs is still not fully understood.

Many biophysical and non-biophysical models have simulated the generation of SWDs leading to an increased knowledge of their underlying mechanisms.^7-21^ However, these models either provided detailed simulated activity only in a selected territory (i.e., cortical or thalamic) or did not test whether their corticothalamic networks could reproduce physiological activities that are known to be generated by these circuits.^22^ Here, we used our corticothalamic model that faithfully simulates EEG waves of natural sleep, i.e., sleep spindles, delta and slow (<0.1 Hz) waves^23^, to investigate whether single abnormal voltage- or transmitter-gated conductances bring about SWDs of ASs. In particular, we show that an increase in the tonic GABA-A inhibition of first order thalamocortical (TC_FO_) neurons, a decrease in cortical phasic GABA-A inhibition, an increase in cortical AMPA receptor function or an increase in the T-type Ca^2+^ conductance of higher order thalamocortical (TC_HO_) neurons generate ∼4 Hz SWDs (as observed in humans with ASs^2,5^) that invariably start in the neocortex.

## METHODS

### Corticothalamic network model

We used our biophysical model of the corticothalamic network (Fig. S1) that faithfully reproduces the typical EEG waves of natural sleep.^23^ Briefly, the model contains a single cortical column with different numbers of excitatory and inhibitory neurons in layers 2/3 to 6, including regular spiking (RS), intrinsically bursting (IB), fast spiking (FS), early firing (EF), repetitive intrinsically bursting (RIB) and network driver (ND) neurons^22^ (Fig. S1). The connections of the different neuronal populations within a layer and between layers are illustrated in Fig. S1. The thalamic territory has a first-order and a higher-order sector with their respective TC neurons^24^ (TC_FO_ and TC_HO_ neurons), each with reciprocal connections with two sectors of the nucleus reticularis thalami (NRT) (NRT_FO_ and NRT_HO_ neurons, respectively) (Fig. S1). The precise network model organization and the numerical values of the connectivity parameters and of the various intrinsic and synaptic conductances of the different neuronal populations are detailed in Martynas and Crunelli (2022).^23^

### Data analyses

Simulation data were analysed and visualised with the help of custom-written Matlab (MathWorks Inc., USA) routines. Raw EEG signal was filtered and cross-correlated as described in Martynas and Crunelli (2022).^23^ SWD Hilbert transform phase was calculated by band-pass filtering raw EEG traces using Butterworth filter with the following parameters: passband and stopband frequencies centered ±2 Hz and ±4 Hz around the SWD oscillation frequency, respectively, and passband ripple and stopband attenuation being 0.5 dB and 65 dB, respectively. The filtered EEG signal was then subjected to Matlab’s hilbert function. Hilbert phase synchronization index (PSI)^25^ was calculated for two filtered signals obtained using the same filtering parameters as above and then smoothed using moving average window of 1 second duration (for additional data analyses see Supplementary Methods in Martynas and Crunelli (2022).^23^

## RESULTS

As shown in the preceding paper^23^, our thalamocortical model is capable of smoothly transitioning between wakefulness (as evident from a low-amplitude, high frequency EEG) and different EEG waves of natural sleep (depending on the input resistance of its constituent neurons) and it does not enter an overly synchronous activity-mode typical of seizures. However, one particular state of the model is prone to generate ictal states, i.e. the transition between sleep and wakefulness. When the model is in this state, different single membrane conductance changes in either cortical or thalamic neurons do lead to an EEG waveform typical of SWDs of ASs, as described below. Notably, all the changes in different conductances that lead to simulated SWDs have a minimal impact on sleep waves (not shown).

### Selective increase in tonic GABA-A inhibition of TC_FO_ neurons generates SWDs

Evidence in mouse and rat AS models have shown that an increased tonic GABA-A inhibition of TC_FO_ neurons (that results from higher thalamic GABA levels^26^) is necessary and sufficient for AS generation.^27-29^ Moreover, higher levels of GABA were found in the thalamus of a child with ASs^30^ and drugs that are known to increase GABA levels, i.e. vigabatrin and tiagabine, can induce or aggravate ASs in humans.^31,32^ Increasing (by 5%) the leak conductance (g_KL_) in TC_FO_ neurons (in order to mimic the increased tonic GABA-A inhibition observed in genetic AS models^27,28^) led to the appearance of SWDs at ∼4 Hz (Fig. 1A_1_,B_1_) (Table S1), a frequency similar to that in humans with ASs.^2,5^ Further increases in g_KL_ did not change the SWD frequency and duration though decreased and then prolonged the interictal period (Fig. 1A_2_,A_3_) (Table S1).

**Figure 1.**
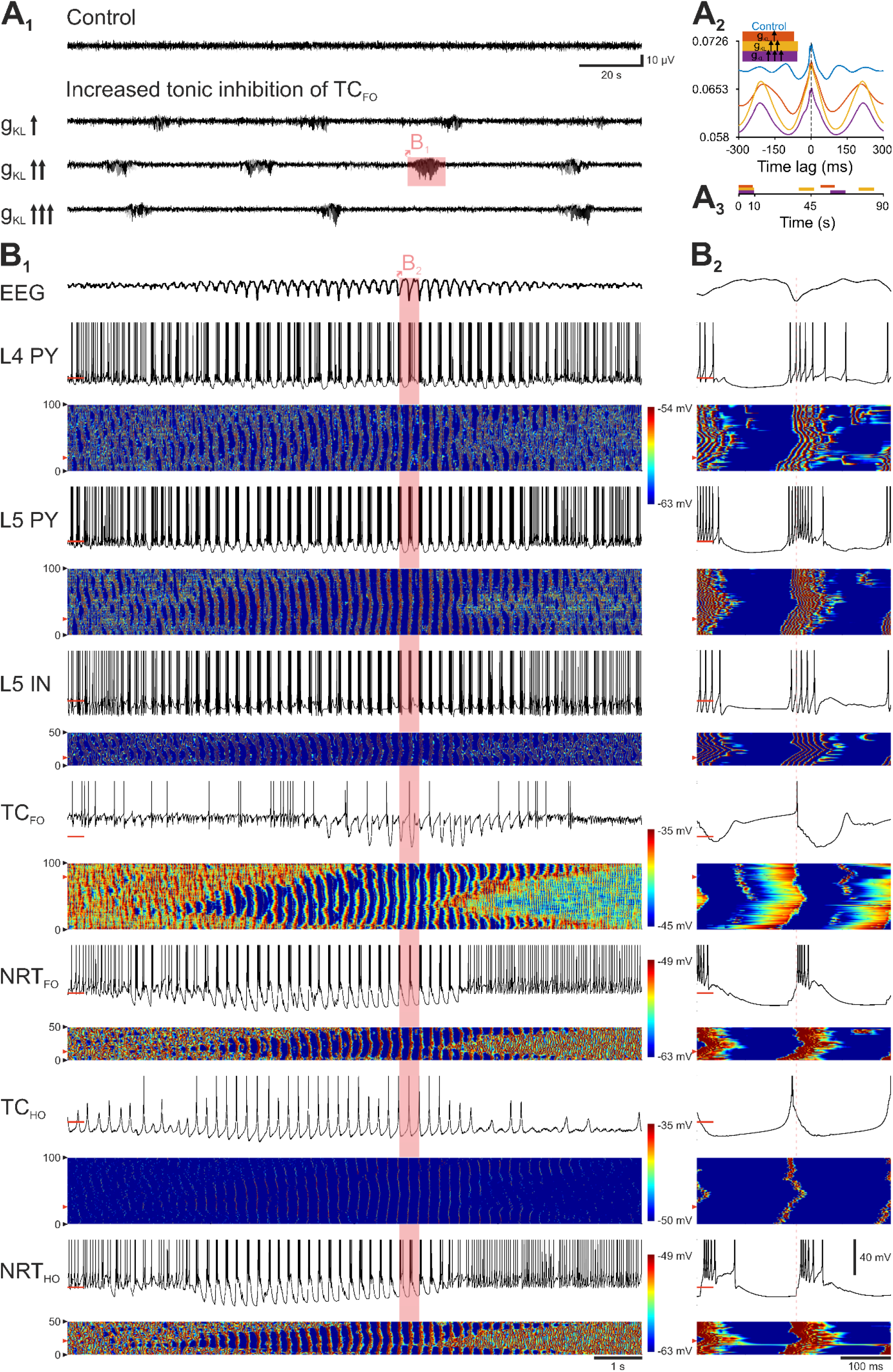
Selective increase in tonic GABA-A inhibition of TC_FO_ neurons elicits SWDs. A_1,_ EEG traces show the induction of spontaneous SWDs at ∼4 Hz after progressive increases in g_KL_ of TC_FO_ neurons, mimicking the constitutively high tonic GABA-A inhibition reported in AS models. The control condition shows simulated desynchronized state typical of relaxed wakefulness. The SWD in the shaded area is expanded in B_1_. A_2_, Cross-correlations between APs of all neurons and the EEG calculated over a 20 min simulation period. Shaded regions represent 95% confidence intervals. A_3_, Schematic timeline showing ictal and interictal periods for different g_KL_ values. Colour-code as in A_2_. B_1_, Top trace: EEG. Panels below show the membrane potential (upper trace) of the indicated neuron and a colour-coded graph of the membrane potential of all neurons of the indicated population. Red bars on the membrane potential traces indicate -60 mV. Red arrowheads in the colour-coded graphs mark the neuron shown in the corresponding membrane potential trace. B_2_, Same as B_1_, shows the expanded SWD cycle highlighted in B1. Vertical red dotted line marks the peak of the SWD spike. L4 PY: pyramidal neuron in cortical layer 4; L5 PY: pyramidal neuron in cortical layer 5; L5 IN: interneuron in cortical layer 5; TC_FO_: first order TC neuron; TC_HO_: higher order TC neuron; NRT_FO_: first order NRT neuron; NRT_HO_: higher order NRT neuron.

At the single cell level, almost all neuronal population increased their total firing during SWDs except TC_FO_ neurons that showed a decrease (Figs. 1B_1_,B_2_ and 2A). The same was observed for ictal burst firing whereas tonic, single action potential (AP) firing decreased (Fig. 2B,C). Indeed, burst firing was the highest contributor to ictal activity in all excitatory and inhibitory cortical neurons (independent from their layer location), but was absent in TC_FO_ neurons and similar to tonic firing in all NRT neurons (Fig. 2D). Notably, all cortical and NRT neurons were never silent during SWDs whereas both TC_FO_ and TC_HO_ neurons were mostly silent ictally or fired tonically (Fig. 2D). When considering the firing dynamics of all ictal APs, all neuronal populations fired at or just after the SWD spike except TC_FO_ and TC_HO_ neurons that fired ∼30 and 20 ms, respectively, prior to the SWD spike (Fig. 2E_1_,F_1_). This is also reflected in the firing phase evolution throughout the SWD with TC_FO_ and TC_HO_ cells showing a positive phase through most of the SWD (leading) while cortical cells showing zero or slightly negative phase over the same period (lagging) (Fig. 2G_1_). However, when only the first AP of an SWD cycle was considered, all neurons fired ∼10 ms before the SWD spike (Fig. 2E_2_), and almost all neuron types had a smaller peak ∼80 ms prior to the SWD spike (Fig. 2F_2_). Notably, further increases in the tonic GABA-A inhibition of TC_FO_ neurons moved the peak of the first AP in each cycle to the left and to the right in TC_FO_ and layer 4 pyramidal (L4/PY) neurons, respectively (Fig. 2E_3_,F_3_). Spike-triggered action potential (STAP) histograms, however, do not decisively show which structures are leading during individual oscillation cycles. The temporal evolution of the phase of the first APs indicate that their phases do not remain stable (Fig. 2G_2_). Indeed, whereas the cortex is leading during the initial few seconds of the SWD (Fig. 2G_3_), the TC_FO_ cells briefly catch up and then gradually fall behind the cortical cells again (Fig. 2G_2_,G_4_).

**Figure 2.**
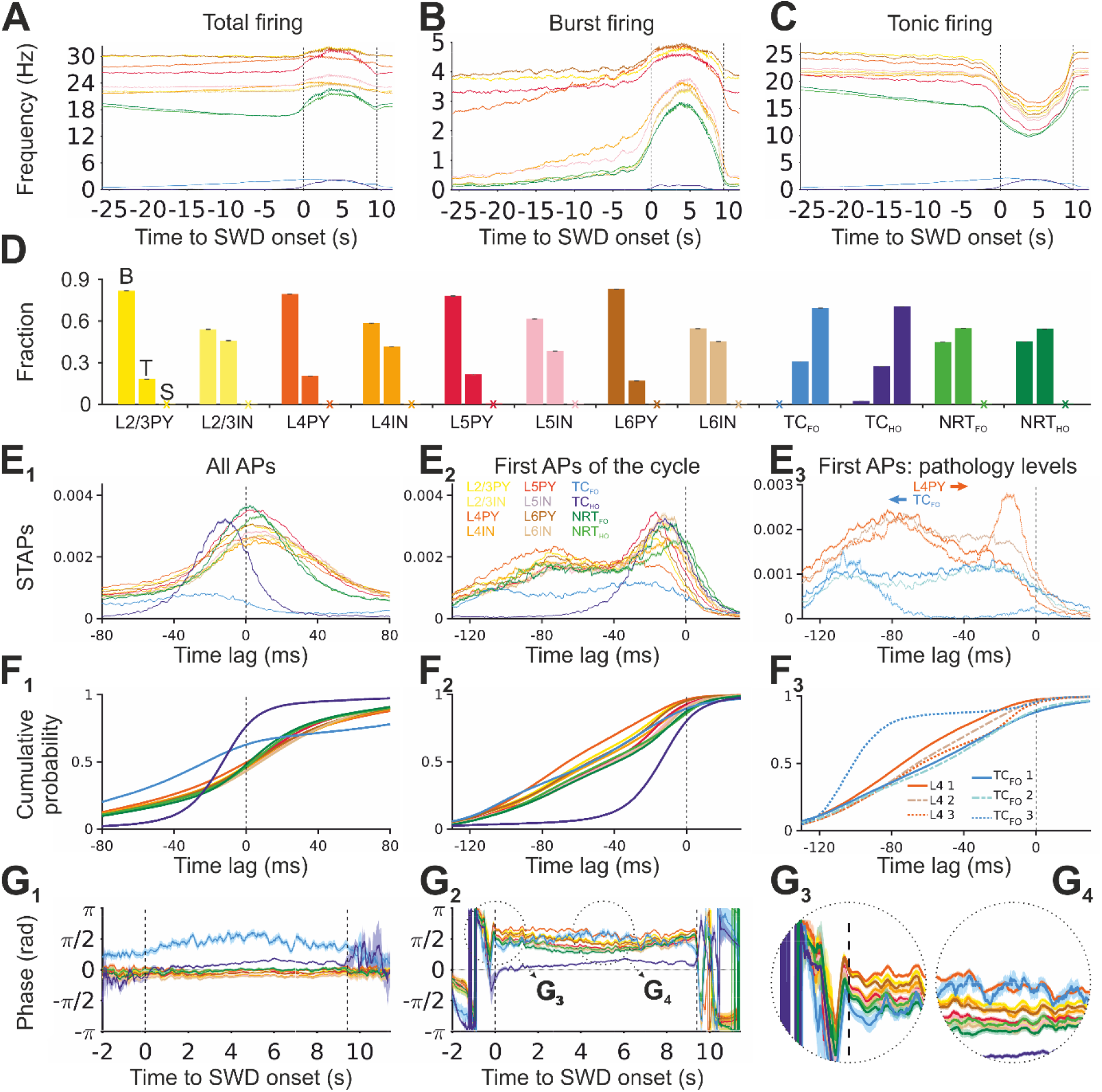
Firing properties during SWDs elicited by increased tonic GABA-A inhibition of TC_FO_ neurons. A-C, Interictal and ictal time-evolution of total, burst and tonic firing frequency for the indicated neuron types. Ictal and interictal periods were linearly scaled to their average durations. The shaded regions represent 95% confidence intervals. Dashed vertical black lines represent the onset and offset of the averaged SWD. Colour-code as in E_2_. D, Mean proportion of the indicated neurons showing burst and tonic firing (B and T column, respectively) and those that are silent (S column). Error bars indicate 95% confidence intervals. E_1_: Cross-correlations between all APs of different neuronal types and the SWD spike (SWD spike triggered action potentials: STAPs). Shaded regions are 95% confidence intervals. Dashed vertical line indicates the peak of the SWD spike. Colour-code as in E_2_. E_2_, Same as E_1_ but only for the first AP in an SWD cycle. E_3_, Same as E_2_, but only for L4PY and TC_FO_ neurons for the three colour-coded g_KL_ values indicated in F_3_ and Fig. 1A_2_. Arrows indicate the shift of the firing peaks as g_KL_ is increased. F_1-3_, Cumulative AP probability corresponding to E_1-3_. G_1_: Hilbert transform mean phase of APs of all cell types with respect to the SWD spike. Different SWDs were linearly scaled to the average duration SWD. Dashed vertical lines indicate the SWD onset and offset. Shaded regions represent 95% confidence intervals. Colour code as in E_2_. G_2_: Same as G_1_ but only for the first AP in an SWD cycle. G_3,4_: Same as G_2_ showing the enlarged regions circled in G_2_. L2/3 PY: pyramidal neuron in cortical layers 2 and 3; L2/3 IN: interneuron in cortical layers 2 and 3; L4 PY: pyramidal neuron in cortical layer 4; L4 IN: interneuron in cortical layer 4; L5 PY: pyramidal neuron in cortical layer 5; L5 IN: interneuron in cortical layer 5; L6 PY: pyramidal neuron in cortical layer 6; L6 IN: interneuron in cortical layer 6; TC_FO_: first order TC neuron; TC_HO_: higher order TC neuron; NRT_FO_: first order NRT neuron; NRT_HO_: higher order NRT neuron.

We then analyzed the temporal dynamics of firing synchrony within and between neuronal populations in the interictal and ictal periods (Fig. 3). Within a given neural type, the stronger progressive increase in synchrony from interictal to ictal periods was observed in NRT_FO_ and NRT_HO_ neurons while the smallest occurred in TC_FO_ and TC_HO_ neurons (Fig. 3A_1,2_). Among different populations, those involving all possible pairs of thalamic neurons showed the highest progressive synchrony as did the layer 5 pyramidal neurons (L5PY) pairs with either NRT or TC neurons, whereas the synchrony between L4PY and TC_HO_ neurons gradually decreased (Fig. 3B_3_). Thus, in summary, the temporal dynamics of increased synchrony progresses from thalamic and thalamocortical-cortical neuron pairs to NRT neuron pairs and then to cortical and NRT neuron pairs (Fig. 3C).

**Figure 3.**
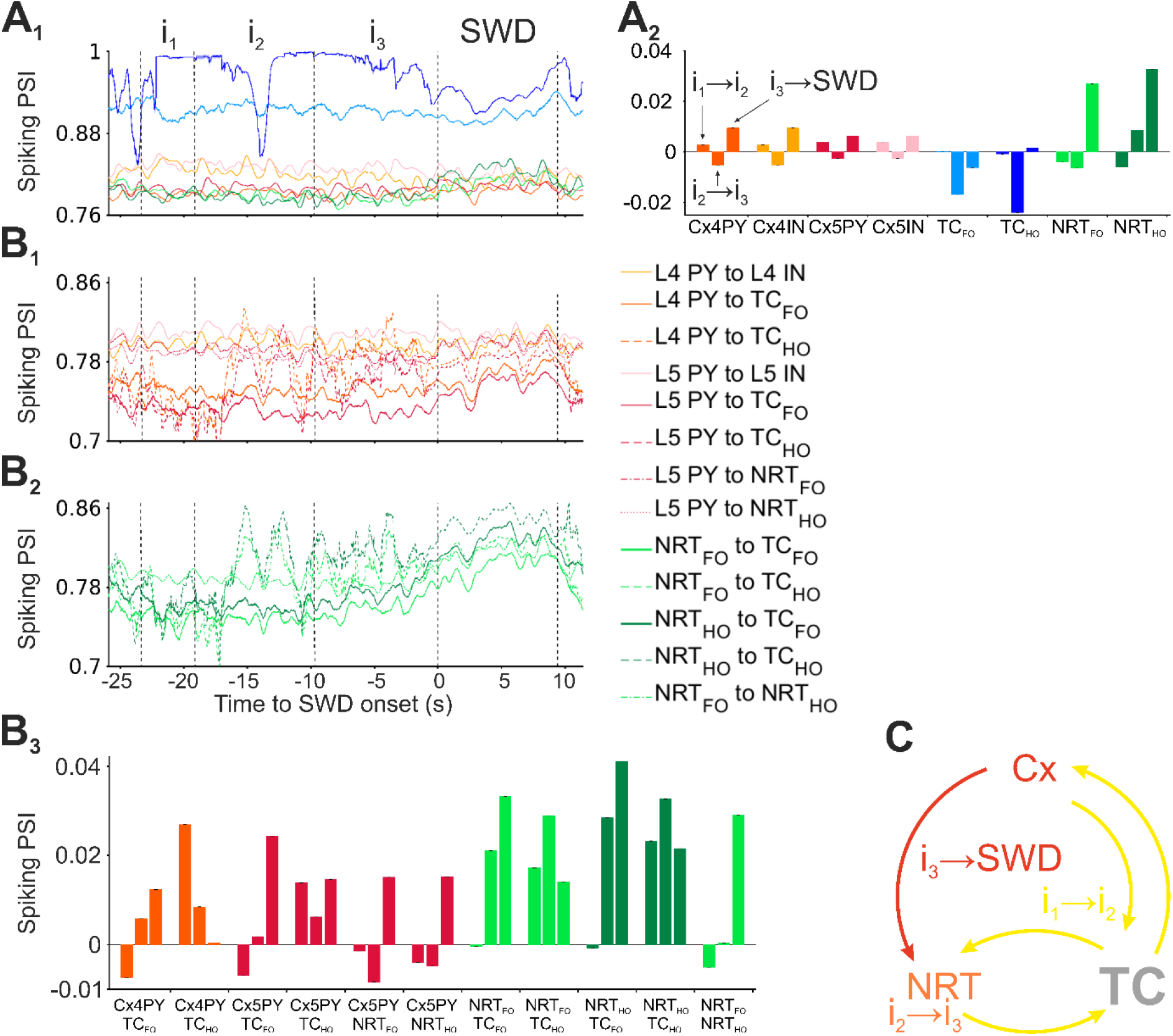
Time-evolution of interictal and ictal firing synchrony for SWDs elicited by increased tonic GABA-A inhibition of TC_FO_ neurons. A_1_, Ictal and interictal mean phase synchronisation index (PSI) of APs within a neuronal population (colour-code as in Fig. 2E2). Ictal and interictal periods were linearly scaled to their average durations. Dashed vertical black lines indicate different parts of ictal and interictal periods: i_1_ marks a section from 1/6 to 1/3 of the interictal period, i_2_ marks the 1/3 to 2/3 section, i_3_ marks the final third of the interictal period, and the last two lines indicate the start and end of the ictal period. A_*2*_, Changes in PSI during interictal and ictal periods. For each indicated neuronal population, the left bar is the PSI change between i_1_ and i_2_ (i_1_ → i_2_), the middle bar between i_2_ and i_3_ (i_2_ → i_3_), and the right bar between i_3_ and the ictal period (i_3_ → SWD). B_1,2_, Evolution of ictal and interictal PSI between different cortical (B_1_) and thalamic (B_2_) populations (colour-code on the right). Vertical dashed black lines demarcate the same regions as in A_1_. B_3,_ Changes in PSI of different neuronal populations over interictal and ictal periods. For each neuronal population pair, the three bars are as in A_2_. C, Schematic representation of the evolution of PSI. Brian areas and their connections shaded in yellow show increases in PSIs between i_1_ and i_2_ and represent the initial synchronization stage (corresponding to bar 1 in all comparison groups of A_2_ and B_3_). Orange and red colours mark PSI increases during the second synchronization stage (between i_2_ and i_3_) and the final synchronization stage (between i_3_ and the ictal period), respectively.

### SWD generation by other selective alterations in inhibitory and excitatory conductances

Selectively decreasing (by 10 and 25%) the conductance of the phasic GABA-A inhibition (g_GABAa_) in all cortical neurons led to SWDs (as observed *in vivo* experiments^33-36^) of progressively larger amplitude and increasingly longer interictal periods (Fig. 4B_1-3_) (Table S1). Compared to the SWDs generated by an increase in thalamic tonic GABA-A inhibition, stronger synchrony was observed between L5PY and all NRT neuron pairs as well as between L4PY and TC_HO_ neuron pairs in the simulated activity induced by a decreased cortical g_GABAa_ (Fig. 4B_4_).

**Figure 4.**
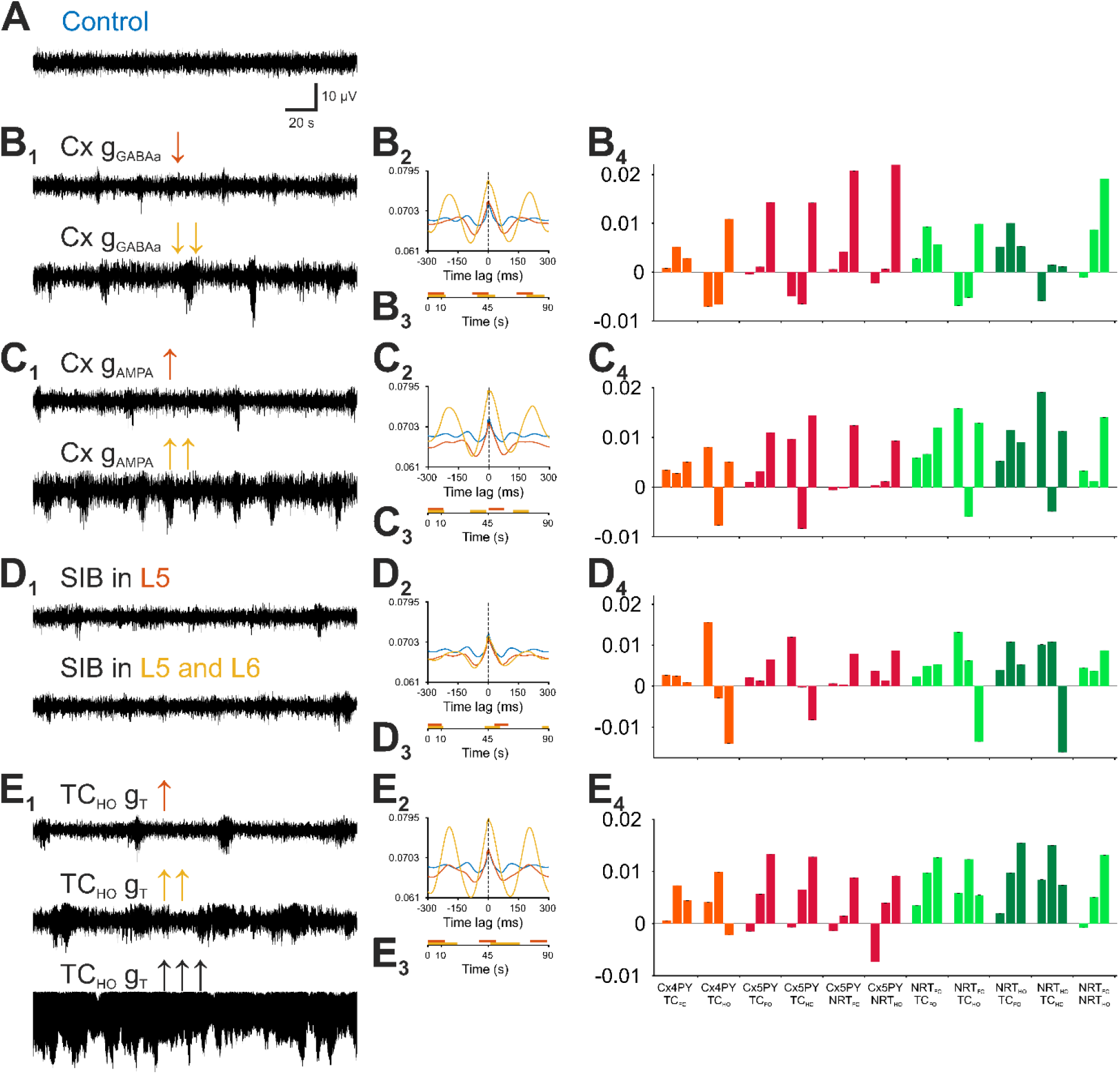
Thalamic and cortical abnormalities can independently induce SWDs. A, EEG showing a period of simulated desynchronised state. B_1_: SWDs elicited by progressive decreases in the GABA-A conductance (g_GABAa_) of all cortical neurons. B_2_, Cross-correlations between APs of all cells and EEG (over a 20 min simulation period) with decreased cortical g_GABAa_. Colour-code as in B1. Shaded regions represent 95% confidence intervals. B_3_, Schematic SWD timeline showing SWD duration and frequency of occurrence for different cortical g_GABAa_. B_4_, Change in the firing PSI between the indicated neuronal populations. For each neuronal population pair, the left bar is the PSI change between i_1_ and i_2_ (i_1_ → i_2_), the middle bar between i_2_ and i_3_ (i_2_ → i_3_), the right bar between i_3_ and the ictal period (i_3_ → SWD), as indicated in Fig. 3A2. C_1-4_, same as B_1-4_ but showing SWDs following increases in the cortical AMPA receptor conductance (g_AMPA_). D_1-4_, same as B_1-4_ but showing SWDs elicited by the addition of strongly intrinsically bursting (SIB) neurons in cortical layer 5 (L5) only or in both L5 and cortical layer 6 (L6). E_1-4_, same as B_1-4_ but showing SWDs after increases in the T-type Ca^2+^ conductance (g_T_) of TC_HO_ cells. Note the absence status reached with the highest increase of g_T_ in these thalamic neurons.

Progressively increasing (by 5 and 40%) the conductance of cortical AMPA receptors (g_AMPA_) also led to more frequent SWDs of increasing amplitude (Fig. 4C_1,3_) (Table S1) and lower synchrony between L5PY and NRT neurons, compared to SWDs elicited by decreased cortical g_GABAa_ (Fig. 4C_4_).

A higher number of cortical strongly intrinsically bursting (SIB) neurons has been reported in genetic rat models of ASs^37,38^: its implementation in the model indeed generated SWDs though of a small amplitude compared to changes in other conductances (Fig. 4D_1-3_) (Table S1) and with a characteristic interictal-to-ictal decrease in synchrony in L4PY-TC_HO_, L5PY-TC_HO_, NRT_HO_-TC_FO_ and NRT_HO_-TC_HO_ neuron pairs (Fig. 4D_4_).

Finally, increasing (by 5 and 10%) the conductance of the T-type Ca^2+^ channels (g_T_) in TC_HO_ neurons generated progressively longer SWDs, ultimately leading to absence status (Fig. 4E_1,3_) (Table S1), and a gradual increase in synchrony among almost all neuronal pairs (Fig. 4E_4_).

### Critical conductances of simulated SWDs

Having established that our model reproduces SWDs elicited by either increasing thalamic tonic or decreasing cortical phasic GABA-A inhibition, we next studied which effect other conductances have on these simulated SWDs. Removing g_T_ from TC_FO_ neurons did not abolish SWDs, as recently reported^39^, but actually increased their amplitude and decreased the interictal period duration (Fig. 5B_1,2_). In contrast, removing g_T_ from TC_HO_ neurons abolished SWDs elicited by both increased thalamic tonic and decreased cortical phasic GABA-A inhibition (Fig. 5C_1,2_) as did g_T_ removal from all types of NRT neurons (Fig. 5D_1,2_).

**Figure 5.**
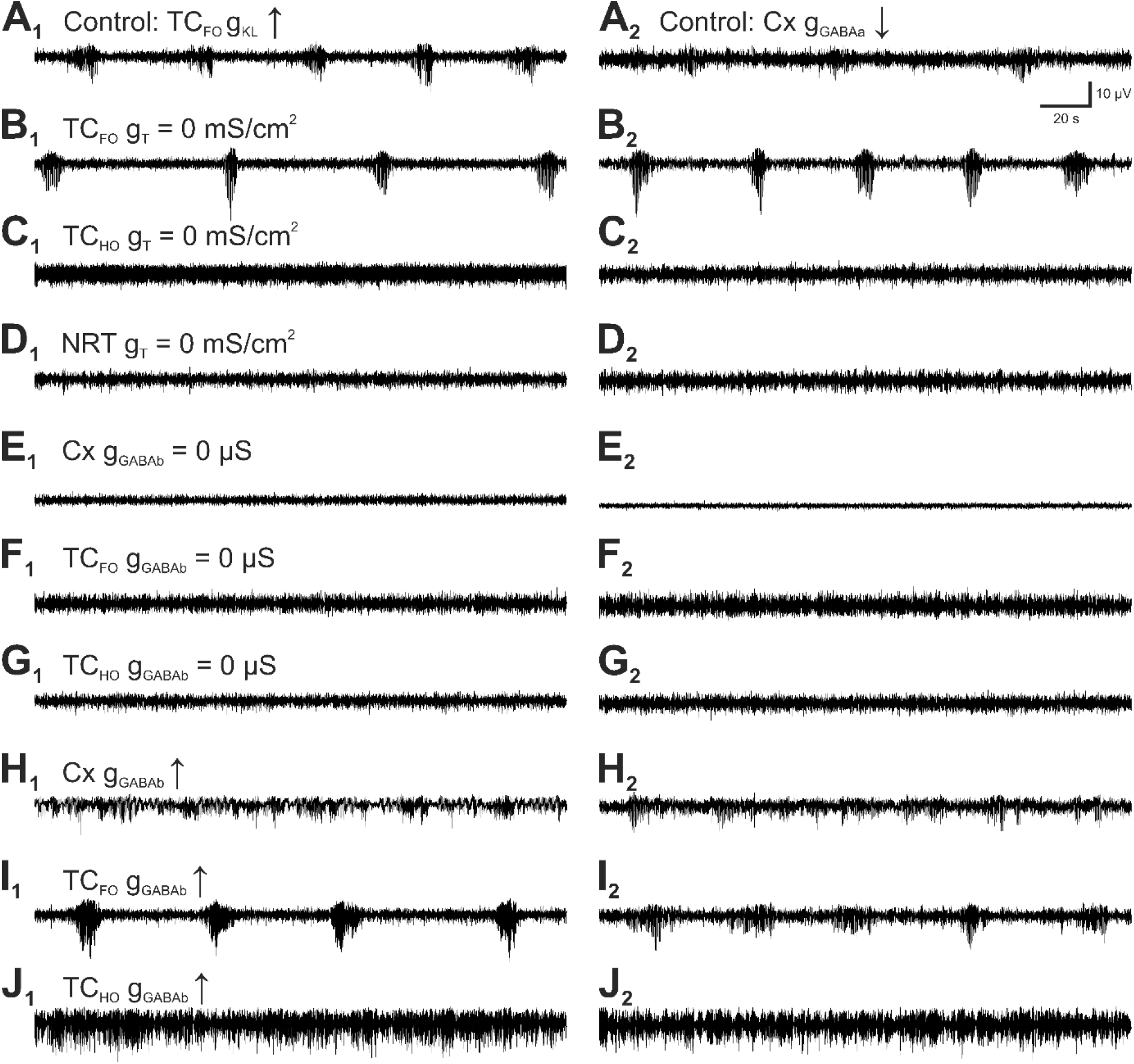
Essential contribution of various voltage- and transmitter-gated conductances to simulated SWDs. A_1_, Control SWDs elicited by increased g_KL_ in TC_FO_ neurons. A_2_: Control SWDs elicited by decreased neocortical g_GABAa_. B_1,2_, SWDs persist after blocking g_T_ in TC_FO_ neurons. C_1,2_, SWDs are not generated when g_T_ is blocked in TC_HO_ neurons. D_1,2_, SWDs are not elicited when g_T_ is blocked in all NRT neurons. E_1,2_, SWDs are blocked after removing g_GABAb_ in all neocortical neurons. F_1,2_, SWDs are blocked after removing g_GABAb_ in TC_FO_ neurons. G_1,2_, SWDs are blocked after removing g_GABAb_ in TC_HO_ neurons. H_1,2_, Smaller amplitude, almost continuous SWDs are elicited when g_GABAb_ is increased in neocortical neurons. I_1,2_, the SWD amplitude and the interictal period are increased when g_GABAb_ is increased in TC_FO_ neurons. J_1,2_, Absence status is generated when g_GABAb_ is increased in TC_HO_ neurons.

Blocking the conductances of GABA-B receptors (g_GABAb_) in either all cortical or thalamic neurons abolished SWDs generated by increased thalamic tonic and decreased cortical phasic GABA-A inhibition (Fig. 5E_1,2_,F_1,2_,G_1,2_) as shown experimentally^40-43^. In contrast, increasing g_GABAb_ in cortical neurons decreased the amplitude of SWDs and markedly increased their duration (Fig. 5H_1,2_). Enhancing g_GABAb_ in TC_FO_ neurons increased the amplitude and the interictal period of SWDs elicited by the increased thalamic tonic GABA-A inhibition (Fig. 5I_1_), whereas it decreased the interictal period of the SWDs simulated by a decreased cortical phasic GABA-A inhibition (Fig. 5I_2_). Finally, increasing g_GABAb_ in TC_HO_ neurons led to absence status in both models (Fig. J_1,2_).

Since the non-selective cation conductance (g_CAN_) plays a key role in some EEG waves of natural^44-46^ and simulated^23^ sleep and its involvement in ASs has not been studied before, we investigated whether it is necessary for simulated SWDs. Removing g_CAN_ from all TC neurons had little effect on SWDs (Fig. 6B_1,2_) whereas its removal from NRT neurons led to absence status (Fig. 6C_1,2_).

**Figure 6.**
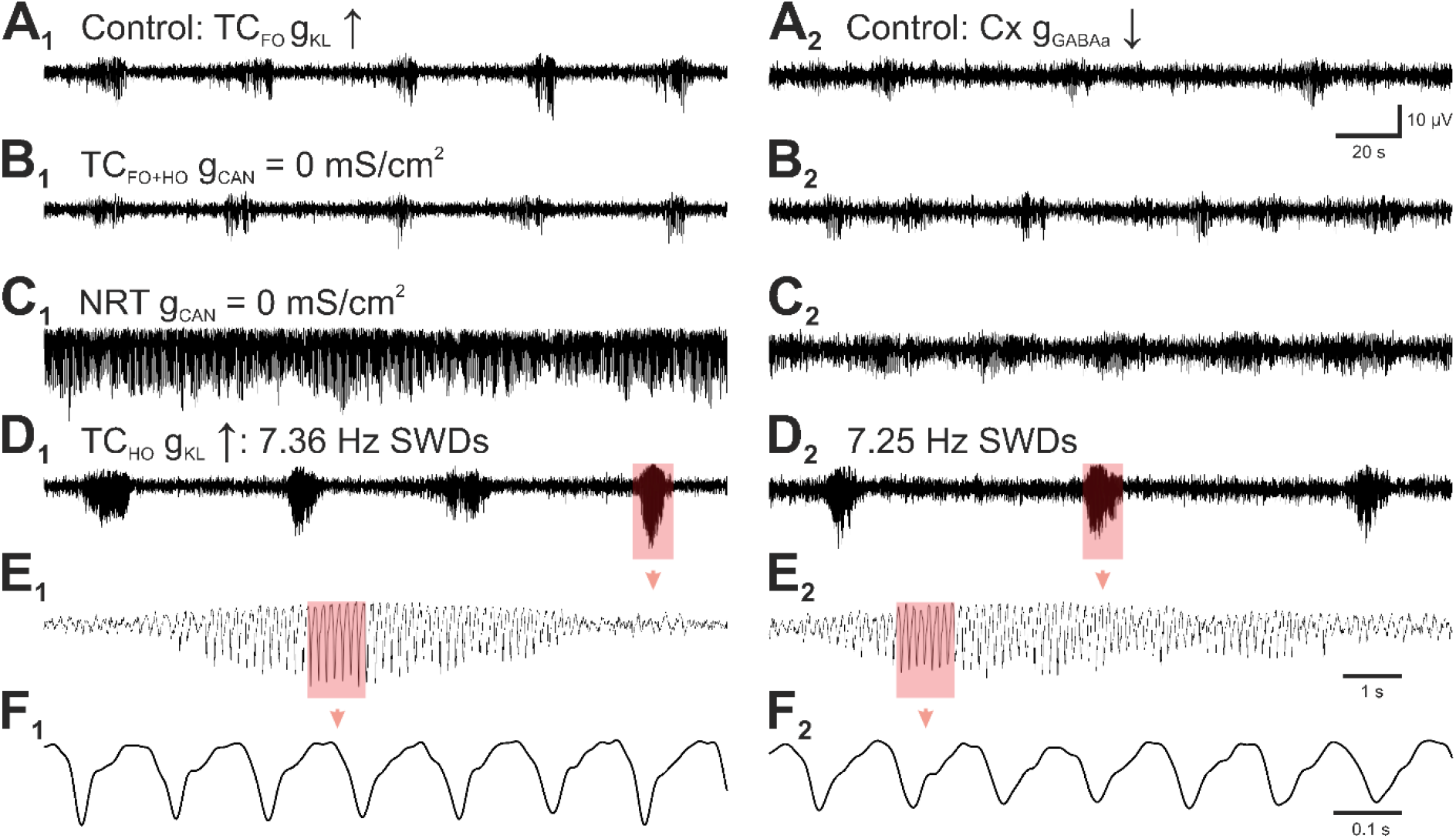
Effect of I_CAN_ and control of SWD frequency by g_KL_ of TC_HO_ neurons. A_1_, Control SWDs elicited by increased g_KL_ in TC_FO_ neurons. A_2_, Control SWDs elicited by decreased neocortical g_GABAa_. B_1_, Blocking the conductance of the non-selective cation current (g_CAN_) in TC_FO_ and TC_HO_ neurons decreases the amplitude of SWDs elicited by the increased g_KL_ in TC_FO_ neurons. B_2_, Blocking g_CAN_ in TC_FO_ and TC_HO_ neurons has very little effect on the SWDs generated by the decreased neocortical g_GABAa_. C_1_, Blocking g_CAN_ in all NRT neurons transform the well separated SWDs elicited by the increased g_KL_ in TC_FO_ neurons into absence status. C_2_, Blocking g_CAN_ in all NRT neurons markedly prolongs the duration of the SWDs elicited by a decreased neocortical g_GABAa_. D_1,2_, Increasing g_KL_ in TC_HO_ neurons leads to SWDs at ∼7 Hz. E_1,2_, Enlargement of the 7 Hz SWDs highlighted in D_1,2_. F_1,2_, Enlargement of the 7 Hz SWD section highlighted in E_1,2_.

Finally, we assessed which conductance was critical for determining the simulated intra-SWD frequency as it represents a major difference between ASs in humans and animal models. We found that a decrease of the leak conductance of TC_HO_ neurons increased the frequency of SWDs from ∼4 Hz to the 7-8 Hz (Fig. 5D_1,2_) that is typical of mouse and rat genetic and pharmacological models.^1,3-5^

## DISCUSSION

The main finding of this study is the ability to faithfully reproduce SWDs at the 4 Hz frequency observed in human ASs by single modifications of neocortical or thalamic conductances in a corticothalamic model that faithfully reproduces the intrinsic and network activity observed in neocortical and thalamic territories during natural sleep.^23^ To the best of our knowledge, this is the most detailed large-scale model dedicated to simulate SWDs, and its component parts and their connectivity patterns were replicated with a high degree of fidelity to experimental data.^22^ Constructing a multi-purpose model guards against an implementation bias of favouring a particular (patho)physiological regime. In fact, no previous attempt at modelling SWDs had this level of physiological validity.

### Model limitations

Notwithstanding, our model has a number of limitations (see Martynas and Crunelli^23^ for details). In the absence of direct measurements, the T-type Ca^2+^ current implemented in various types of neocortical neurons was also guided by the ability of these neurons to faithfully reproduced intrinsic slow (<1 Hz) waves.^23^ Moreover, though no detailed parameters exist for the persistent Na^+^ current in NRT neurons, this current (with biophysical properties similar to those reported for TC neurons^44^) had to be introduced in NRT neurons in order to faithfully reproduced the intrinsic slow (<1 Hz) waves observed in *in vitro* studies.^46^ Finally, since no voltage-clamp study has been performed in higher-order thalamic nuclei, the biophysical properties of the conductances of TC_HO_ neurons were inferred from current-clamp data^47,48^ and/or adapted from TC_FO_ neurons.^49,50^

### Simulation strength

The solidity of our simulated SWDs is supported by two major findings. First, our model faithfully reproduces the three main EEG waves generated by corticothalamic networks during sleep, i.e., spindle, delta and slow (>1 Hz) waves^23^, and these natural rhythms are only minimally affected by the different changes in single voltage- and transmitter-gated conductances that lead to SWDs. Second, our model is capable of reproducing many experimental findings after implementing the different abnormalities that are known to be present in humans with, and genetic models of, ASs.

In particular, our model generates ∼4 Hz SWDs following:

1. blockade of neocortical phasic GABA-A inhibition, as shown experimentally following intracortical injection of the weak and potent GABA-A antagonists penicillin and bicuculline, respectively^33-36,51-53^;
2. increasing the tonic GABA-A inhibition of TC_FO_ neurons, as shown in different genetic models of ASs^27^, i.e. the GAERS (Genetic Absence Epilepsy Rats from Strasbourg) rats and the stargazer and lethargic mouse models;
3. enhancing GABA-B inhibition in either thalamic and cortical territory, as shown by the generation and aggravation of SWDs in normal mice and rats and in genetic AS models, respectively, following systemic, intracortical and intrathalamic injection of GABA-B receptor agonists^40-43^;
4. increasing the T-type Ca^2+^ channel function in TC_HO_ neurons, as reported by Gorji et al. (2011)^47^ and Seidenbecher et al. (2001)^48^;
5. enhancing the number of intrinsically bursting cells in layers 5/6, as observed in the Wistar Albino Glaxo Rats from Rijswijk^37^ and the GAERS genetic models of ASs.^38^

Our simulations also show that SWDs are abolished or reduced following 1) blockade of cortical or thalamic GABA-B receptors as observed in different genetic and pharmacological models of ASs following systemic, intracortical or intrathalamic injection of selective GABA-B receptor antagonists^40-43,^; and 2) removal of T-type Ca^2+^ channels in NRT neurons, as seen following intra-NRT infusion of TTA-P2^39^, a potent and selective blocker of these channels^54^, in GAERS rats. In contrast, simulated SWDs are unaffected by blocking T-type Ca^2+^ channels in TC_FO_ neurons as reported by McCafferty et al. (2018).^39^

Finally, the strength of our results is also supported by their similarities with the following experimental findings:

1. TC_FO_ neurons, as those in the ventrobasal thalamus, are mostly silent during SWDs^39,55^;
2. burst firing of NRT and cortical neurons increases during SWDs^39,56^;
3. tonic firing is reduced in all types of cells, as shown experiemtallty^39^, except in TC_HO_ cells for which no data are available at present;
4. the burst firing of cortical pyramidal neurons and interneurons in all layers is increased ictally compared to interical periods^38,39,57^;
5. the transition between sleep and quiet wakefulness is the vigilance states where most SWDs occur.^58,59^

### Predictions from simulated SWDs

A number of key testable predictions originates from the results of our study:

1. T-type Ca^2+^ channel-mediated burst firing in TC_HO_ neurons is necessary to elicit SWDs;
2. depolarization of TC_FO_ neurons prevents SWD generation or interfere with ongoing SWDs;
3. I_CAN_ of NRT neurons is essential for termination of SWDs;
4. g_KL_ of TC_HO_ neurons is a key determinant of SWD frequency;
5. absence status occurs when i) blocking I_CAN_ in NRT neurons, ii) strongly increasing g_T_ in TC_HO_ neurons, and iii) increasing g_GABAb_ in TC_HO_ neurons. These results provide the first mechanistic insight into absence status and have a strong translational significance since both T-type Ca^2+^ channel blockers and GABA-B receptor antagonists are being trialed in human cohorts.^5^

## Acknowledgments

This work was supported by an MRC PhD studentship to MD and by the Ester Floridia Neuroscience Research Foundation (grant 1502 to VC).

## Notes

**Conflict of interest** The authors declare no conflict of interest.

### Competing Interest Statement

The authors have declared no competing interest.

