## Supplementary Information for "Spike-and-wave discharges of absence seizures in a sleep waves-constrained corticothalamic model"

Supplementary Figure 1 page 2

Supplementary Table 1 page 4

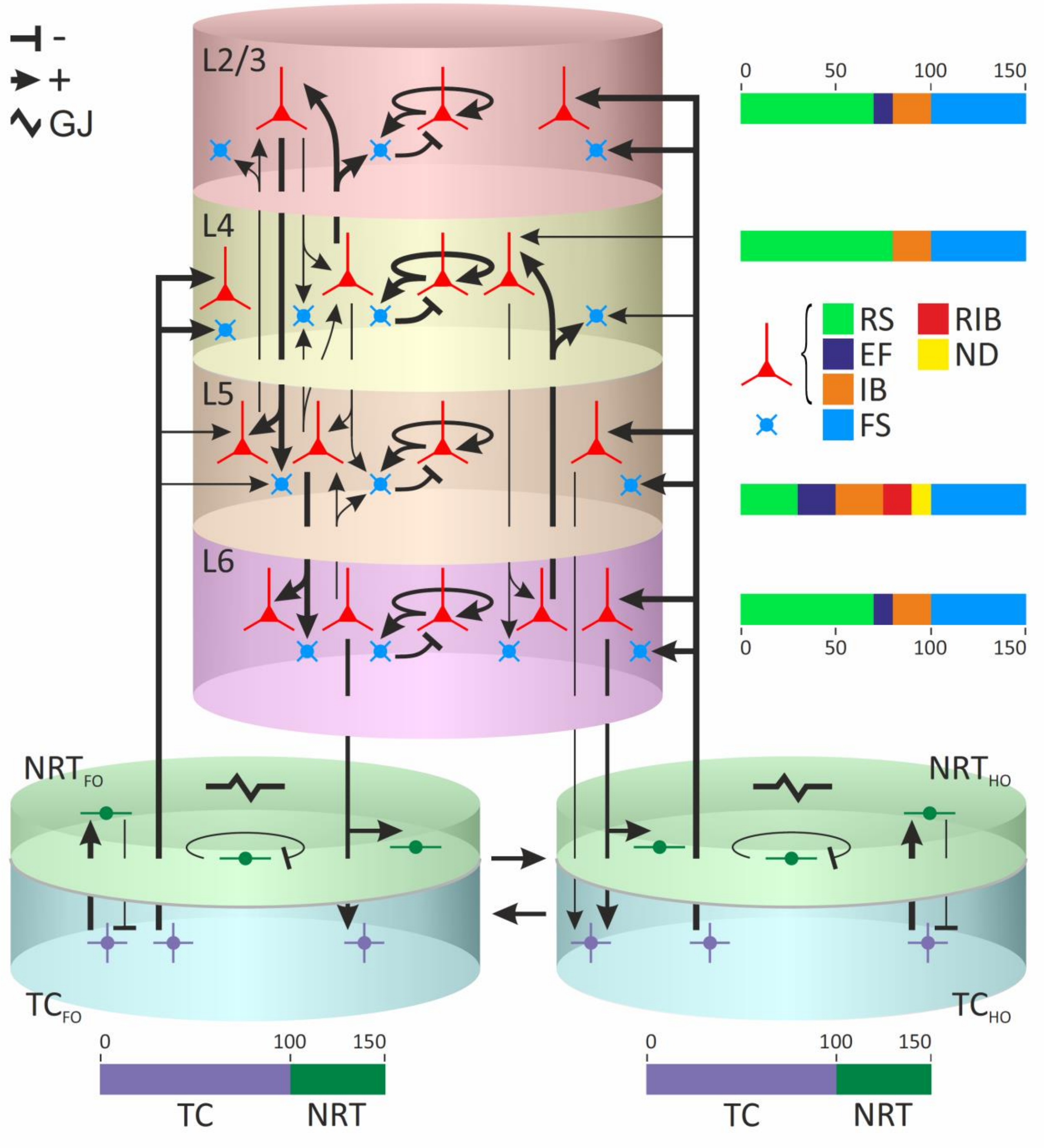


**Figure S1. Corticothalamic model architecture.**

The corticothalamic model consists of 900 neurons distributed in distinctly coloured layers of a single cortical column and two coloured-coded thalamic sectors, a first- and a higher-order sector. Each cortical layer contains of 100 excitatory neurons (detailed neuronal populations are in the colour-coded bars on the right) and 50 fast-spiking (FS) inhibitory neurons. Thalamic cylinders (of both first-order and higher-order sectors) contain 100 thalamocortical (TC) neurons and 50 nucleus reticularis thalami (NRT) neurons. Sharp and blunt arrows represent excitatory and inhibitory synaptic connections, respectively, with the line thickness indicating the synaptic connection strength (see actual values in Supplementary Table 1). The lightning symbol indicates gap junctions that are present only between NRT neurons.

**L2/3**: cortical layer 2/3

**L4**: cortical layer 4

**L5**: cortical layer 5

**L6**: cortical layer 6

**RS**: regular spiking neuron

**EF**: early firing neuron

**IB**: intrinsically bursting neuron

**RIB**: repetitive intrinsically bursting neuron

**ND**: network driver neuron

**FS**: fast spiking neuron

**$TC_{FO}$**: thalamocortical neurons of first-order thalamic nucleus

**$TC_{HO}$**: thalamocortical neurons of higher-order thalamic nucleus

**$NRT_{FO}$**: nucleus reticularis thalami neurons connected to first-order nucleus,

**$NRT_{HO}$**: nucleus reticularis thalami neurons connected to higher-order nucleus.

**Table S1. Parameters of SWDs induced by different pathologies.**

| Pathology type | Tonic inhibition of $TC_{FO}$ cells | | | Reduced cortical $GABA_A$ conductance | | Increased cortical AMPA conductance | | SIB in the cortex | | Increased $I_T$ in $TC_{HO}$ | |
|---|---|---|---|---|---|---|---|---|---|---|---|
| Pathology level | 1 | 2 | 3 | 1 | 2 | 1 | 2 | SIB in L5 | SIB in L5/6 | 1 | 2 |
| SWD incidence ($min^{-1}$) | 1.16 | 1.57 | 1.01 | 1.82 | 1.62 | 1.32 | 1.85 | 1.16 | 1.37 | 1.57 | 1.32 |
| Mean SWD duration (s) | 8.76 ± 0.29 | 9.43 ± 0.19 | 9.32 ± 0.25 | 11.84 ± 0.43 | 13.19 ± 0.41 | 11.52 ± 0.46 | 11.85 ± 0.23 | 10.49 ± 0.62 | 11.21 ± 0.4 | 12.47 ± 0.39 | 21.96 ± 0.77 |
| Mean interictal period (s) | 42.08 ± 1.35 | 27.93 ± 0.71 | 47.71 ± 1.68 | 21.3 ± 0.75 | 23.58 ± 0.84 | 33.62 ± 1.71 | 19.81 ± 0.8 | 39.01 ± 2.3 | 31.38 ± 1.33 | 25.64 ± 1.01 | 24.62 ± 1.35 |
| Time spent in SWDs (%) | 17.23 | 25.23 | 16.34 | 35.72 | 35.87 | 25.53 | 37.44 | 21.19 | 26.32 | 32.72 | 47.15 |
| Mean spike-to-wave amplitude (µV) | 5.83 ± 0.0057 | 7.12 ± 0.005 | 6.78 ± 0.007 | 4.44 ± 0.002 | 6.85 ± 0.003 | 4.72 ± 0.003 | 8.24 ± 0.004 | 3.86 ± 0.004 | 4.25 ± 0.003 | 4.93 ± 0.003 | 8.45 ± 0.002 |
| Mean PSI | 0.64 ± 0.00002 | 0.7 ± 0.00001 | 0.69 ± 0.00002 | 0.64 ± 0.00001 | 0.52 ± 0.000009 | 0.57 ± 0.00001 | 0.65 ± 0.00001 | 0.56 ± 0.00002 | 0.59 ±0.00001 | 0.58 ± 0.00001 | 0.82 ± 0.00001 |
| AP cross-correlation amplitude | 0.02 | 0.026 | 0.019 | 0.019 | 0.043 | 0.019 | 0.051 | 0.015 | 0.018 | 0.019 | 0.059 |
| AP-EEG correlation amplitude | 0.0078 | 0.0098 | 0.0071 | 0.0077 | 0.014 | 0.0078 | 0.015 | 0.0062 | 0.0073 | 0.0075 | 0.018 |
| Intra-SWD freq. (Hz) | 4.79 | 4.75 | 4.66 | 4.79 | 4.6 | 4.86 | 4.64 | 5.06 | 5.19 | 5.09 | 4.99 |
| Slowing down of intra-SWD freq. (yes/stable/increase) | Yes | Yes | Increase | Yes | Increase | Yes | Increase | Stable | Increase | Stable | Stable |
| Presence of poly-spikes (yes/no) | Yes | Yes | Yes | Yes | Yes | Yes | Yes | Yes | Yes | Yes | Yes |
| Leading structure | L4 | L4 | $TC_{FO}$ | L4 | L4 | L4 | L4 | L4 | L4 | L4 | L4 |
| Mean θ of $1^{st}$ L2/3 AP in $1^{st}$ sec of SWD (rad) | 1.98 ± 0.052 | 1.94 ± 0.05 | 1.97 ± 0.057 | 1.77 ± 0.05 | 2.37 ± 0.19 | 1.91 ± 0.05 | 1.29 ± 0.05 | 1.98 ± 0.054 | 1.9 ± 0.051 | 1.97 ± 0.045 | 1.55 ± 0.066 |
| Mean θ of $1^{st}$ L4 AP in $1^{st}$ sec of SWD (rad) | 2.13 ± 0.041 | 2.045 ± 0.042 | 2.01 ± 0.054 | 2.29 ± 0.03 | 2.76 ± 0.11 | 2.17 ± 0.046 | 2 ± 0.035 | 2.19 ± 0.044 | 2.15 ± 0.041 | 2.18 ± 0.037 | 2.02 ± 0.043 |

| | | | | | | | | | | | |
|---|---|---|---|---|---|---|---|---|---|---|---|
| Mean θ of 1st L5 AP in 1st sec of SWD (rad) | 1.69 ± 0.062 | 1.61 ± 0.059 | 1.66 ± 0.063 | 1.45 ± 0.059 | 2.06 ± 0.26 | 1.66 ± 0.065 | 1.04 ± 0.051 | 1.72 ± 0.066 | 1.56 ± 0.064 | 1.74 ± 0.053 | 1.13 ± 0.078 |
| Mean θ of 1st L6 AP in 1st sec of SWD (rad) | 1.91 ± 0.055 | 1.87 ± 0.053 | 1.95 ± 0.059 | 1.7 ± 0.055 | 2.3 ± 0.22 | 1.86 ± 0.06 | 1.16 ± 0.05 | 1.93 ± 0.06 | 1.81 ± 0.059 | 1.91 ± 0.048 | 1.42 ± 0.075 |
| Mean θ of 1st $NRT_{FO}$ AP in 1st sec of SWD (rad) | 1.59 ± 0.09 | 1.51 ± 0.082 | 1.31 ± 0.11 | 1.92 ± 0.055 | 2.4 ± 0.31 | 1.7 ± 0.09 | 1.69 ± 0.059 | 1.72 ± 0.088 | 1.71 ± 0.079 | 1.75 ± 0.072 | 1.56 ± 0.088 |
| Mean θ of 1st $NRT_{HO}$ AP in 1st sec of SWD (rad) | 1.41 ± 0.108 | 1.37 ± 0.093 | 1.31 ± 0.099 | 1.69 ± 0.072 | 1.95 ± 0.47 | 1.42 ± 0.11 | 1.45 ± 0.073 | 1.4 ± 0.12 | 1.42 ± 0.11 | 1.48 ± 0.096 | 1.05 ± 0.11 |
| Mean θ of 1st $TC_{FO}$ AP in 1st sec of SWD (rad) | 1.51 ± 0.134 | 1.25 ± 0.15 | 1.35 ± 0.33 | 1.88 ± 0.075 | 2.35 ± 0.39 | 1.7 ± 0.13 | 1.76 ± 0.07 | 1.68 ± 0.11 | 1.59 ± 0.1 | 1.74 ± 0.094 | 1.87 ± 0.054 |
| Mean θ of 1st $TC_{HO}$ AP in 1st sec of SWD (rad) | -0.23 ± 0.137 | -0.017 ± 0.104 | 0.4 ± 0.072 | 0.23 ± 0.13 | -0.09 ± 0.33 | -0.19 ± 0.16 | 0.15 ± 0.12 | -0.45 ± 0.16 | -0.41 ± 0.16 | -0.34 ± 0.13 | -0.054 ± 0.089 |
| Fraction of bursting in $TC_{FO}$ cells during SWDs (%) | 0 ± 0 | 0 ± 0 | 0.0034 ± 0.00039 | 0 ± 0 | 0 ± 0 | 0 ± 0 | 0 ± 0 | 0 ± 0 | 0 ± 0 | 0 ± 0 | 0 ± 0 |
| Fraction of tonic spiking in $TC_{FO}$ cells during SWDs (%) | 42.87 ± 0.26 | 30.62 ± 0.22 | 5.43 ± 0.038 | 63.4 ± 0.25 | 54.16 ± 0.31 | 60.03 ± 0.25 | 61.12 ± 0.26 | 61.02 ± 0.25 | 58.01 ± 0.26 | 57.24 ± 0.26 | 56.9 ± 0.26 |

Abbreviations: PSI, phase synchronisation index; freq, frequency; θ, phase.
